# 3D-multiple immunoimaging using whole male organs in rice

**DOI:** 10.1101/2022.05.16.492221

**Authors:** Saori Araki, Hinako Tamotsu, Reina Komiya

## Abstract

The spatiotemporal regulation of proteins and RNAs is essential for the precise development of reproductive tissues in many organisms. The anther, a significant part of the male reproductive organ in plants, contains the somatic cell layers named the anther wall and germ cells located inside. The cell-to-cell communication between the soma and germ cells in the development of male and female tissues is reported, although the mechanism remains unknown in plants. Here, we successfully developed a simple 3D-organ-immunoimaging technique, which distinguishes each single cell from the four somatic cell layers and germ cells without the need for transformation, embedding, sectioning, or clearing in rice. The 3D-immunostaining method is also applicable to the intracellular localization of meiosis-specific proteins in meiocytes—namely MEL1, a germ cell-specific Argonaute in the cytoplasm, and ZEP1, a pachytene marker on meiotic chromosomes. Our study suggests that the 3D-multiple immunostaining method with single-cell and intracellular resolution will contribute to the organ-level elucidation of comprehensive molecular mechanisms and non-cell-autonomous systems.

## Introduction

The development of 3D-organ imaging at a subcellular and single-cellular resolution is essential for a comprehensive understanding of molecular mechanisms because the precise developmental mechanisms in many organisms involve spatiotemporal regulation and cell-to-cell interaction. Argonaute (AGO)-interacting small RNAs form the core machinery of RNA silencing. Several AGO-small RNA complexes spatiotemporally regulate germline development via silencing in higher organisms ^1–4^. The anther is a major part of the male reproductive organ in plants. Anthers consist of germ cells, also known as the pollen and somatic anther walls, which are the three to four outer layers surrounding the germ cells located inside the anthers (Fig. 1A). Anther development is approximately classified into four stages, the premeiosis, early meiosis, meiotic division, and microspore (Supplementary Table S1) ^5^. The development of the somatic cell layer synchronizes with that of the germ cells, suggesting cell-to-cell communication between the soma and germ in anther development ^6,7^. In addition, *Arabidopsis* AGO9 interacting repeat-associated small RNAs regulates the number of the megaspore mother cells ^8^. However, the non-cell-autonomous system is poorly understood in the reproductive tissues of plants. Hence, the development of immunohistochemical technologies using whole tissues can also promote the elucidation of cell-to-cell interactions distinguishable at the single-cell level.

**Figure 1.**
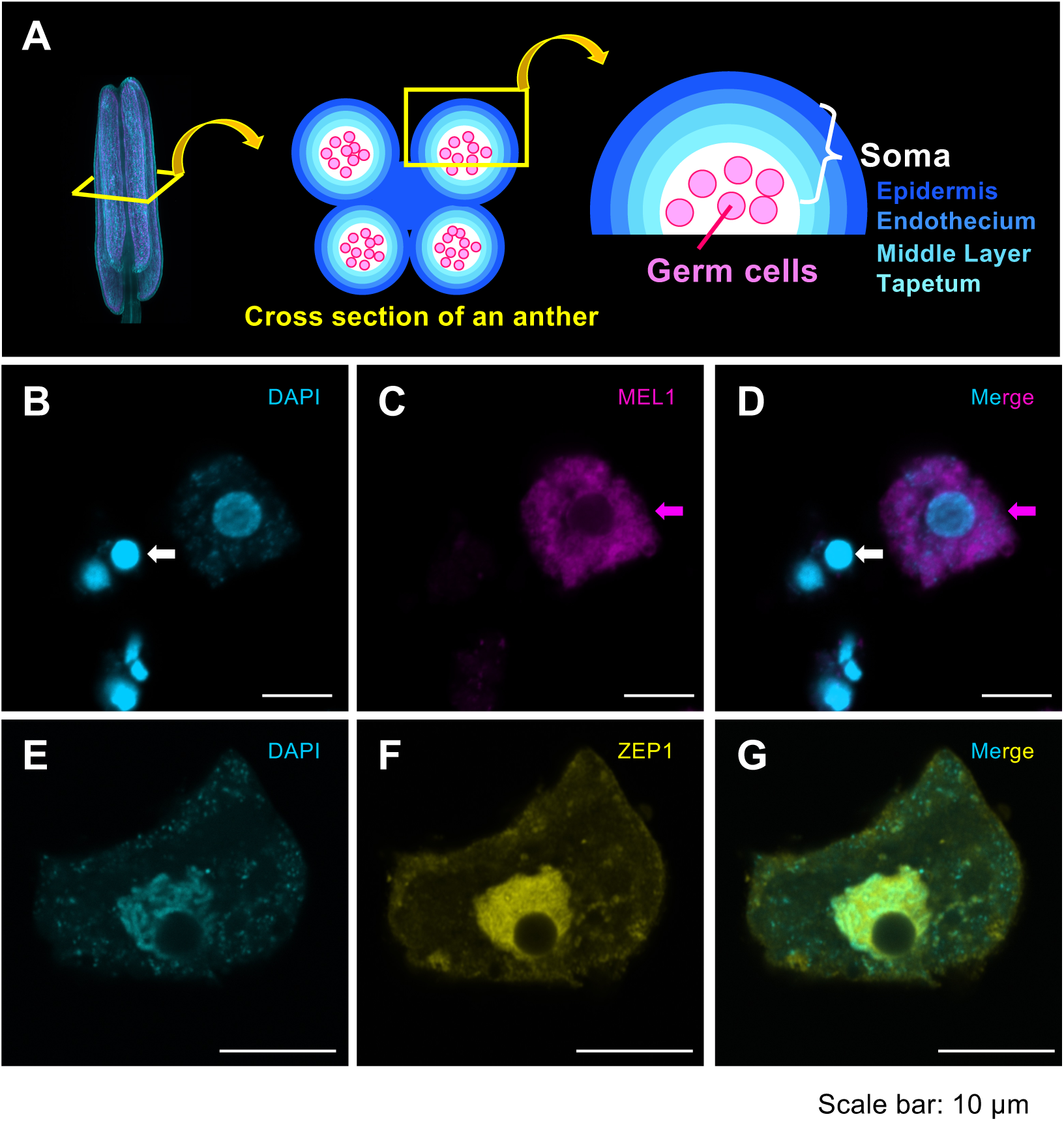
Immunostaining using pollen mother cells for anti-MEL1 and meiocytes for anti-ZEP1. **(A)** Schematic diagram of an anther. The image on the left shows an anther stained with SCRI Renaissance 2200. Anther consists of four locules (middle), and a cross-section of a single locule demonstrates the four somatic anther walls showing the epidermis (Ep), endothecium (En), middle layer (Ml), and tapetum (Ta) and germ cells. (**B-D**) MEL1 localization is restricted in the cytoplasm (magenta). White arrows indicate single somatic cells from four anther walls, although the origins from somatic cell types, Ep, En, Ml, and Ta remain unknown. Magenta arrows indicate the pollen mother germ cell at the premeiotic stage. (**E-G**) The indirect fluorescence signals of ZEP1 were enriched on the meiotic chromosome (yellow). Cyan signals indicate DAPI staining (**B** and **E**). Merged images of B and C, and F and G, respectively (**D** and **G**). Scale bars, 10 μm.

To investigate the structure of tissues and distinguish each single cell, morphological imaging is beneficial in tandem with histochemical staining. However, morphological analysis needs many steps, including the fixation of anthers, embedding in resin, sectioning, and staining ^9,10^. Our previous study of histochemical imaging recently enabled the visualization of the three-dimensional (3D) structure of entire rice anthers using Lightsheet microscopy ^11^. This method can also be used to distinguish the internal structure of the anthers, indicating that morphological analysis is suitable for detecting developmental defects in mutants.

In order to investigate the localization of proteins and elucidate the molecular mechanism, imaging using fluorescent proteins is more established than immunostaining in plants ^12 13^. However, this method requires an additional transformation step to generate plants with the target protein fused to a fluorescent protein. The immunohistochemical method, which can detect epigenetic marker proteins in leaves, is reported to be accompanied by clearing steps ^14^. Using meiocytes which are germ cells during meiosis, the cellulase treatment disrupts the anther wall, thus impeding the distinguishability of which somatic layers the single cells are derived from ^15^. Accordingly, 3D-histological immunoimaging using whole anthers is an essential tool for determining the spatiotemporal regulation and cell-to-cell interaction during plant reproduction. Meiosis is an essential phenomenon in which genetic information is inherited by the subsequent generation. ZEP1 is a component of transverse filaments of the rice synaptonemal complex ^16^. MEIOSIS ARRESTED AT LEPTOTENE1 (MEL1) interacting with 21-nucleotide small RNAs, are required for meiotic progression during early meiosis in rice ^15,17^. *mel1* mutants exhibit defects of ZEP1 elongation on meiotic chromosomes. However, how the MEL1-small RNAs complex causes meiotic silencing via ZEP1 regulation between the nucleolus and cytoplasm in meiocytes remains unknown. Here, we successfully developed a 3D immunostaining method using whole rice anthers to visualize intracellular protein localization, the cytoplasmic MEL1 and filamentous signals of ZEP1, and structure of each single cell in the four somatic cell layers. The advantage of our simple method is that it can distinguish the subcellular localizations of multiple proteins in the four somatic cell layers and germ cells simultaneously, by eliminating the need for transformation, embedding, cross-sectioning and clearing.

## Results

### Creation and evaluation of antibodies

First, we generated the antibodies of MEL1 and ZEP1 as germ-cell markers and a pachytene markers, respectively, to develop the 3D-immunoimaging method using whole rice anthers (see below the material and methods). Automated western analysis using Wes, with MEL1 and ZEP1 antibodies revealed that MEL1 and ZEP1, with the predicted molecular sizes, were expressed at the meiotic stage, where the anthers are 0.6–0.7 mm in size (Supplementary Fig. S1). Moreover, we performed immunostaining using a previously reported method ^10,15^ and detected these antibodies using pollen mother cells for MEL1 and meiocytes for ZEP1. MEL1 was enriched at the cytoplasm in pollen mother cells at premeiosis (Fig. 1B-D). ZEP1 was restricted to the meiotic chromosome at the progression of homologous chromosome synapsis during early meiosis, as previously reported (Fig. 1E-G) ^15,16^. Therefore, these MEL1/ZEP1 antibodies are suitable meiocytes/meiotic landmarks in 3D-multiple immunostaining.

### Immunostaining using whole anthers

Split anthers and degassing prior to immunization with primary and secondary antibodies are indispensable steps in 3D-multiple immunoimaging (Fig. 2A). The anther sizes coincide with the anther-development stages, corresponding to the flower sizes (Table S1). We isolated 0.6–0.7 mm anthers from 4% paraformaldehyde-fixed inflorescences during early meiosis, wherein homologous chromosome synapsis occurs. The four somatic layers of the anther wall, which contains the epidermis (Ep), endothecium (En), middle layer (Ml), and tapetum (Ta), are formed, surrounding the meiocytes at the early meiosis stage (Fig. 1A). For the antibodies to penetrate the whole anthers, the 4% paraformaldehyde-fixed anthers were transferred to distilled water on MAS-coated microscope slides and split into two using a scalpel under a stereomicroscope, taking care to not collapse the structure of the anthers (Fig. 2B-D). This process of anther split contributes to the success of 3D-multiple immunoimaging.

**Figure 2.**
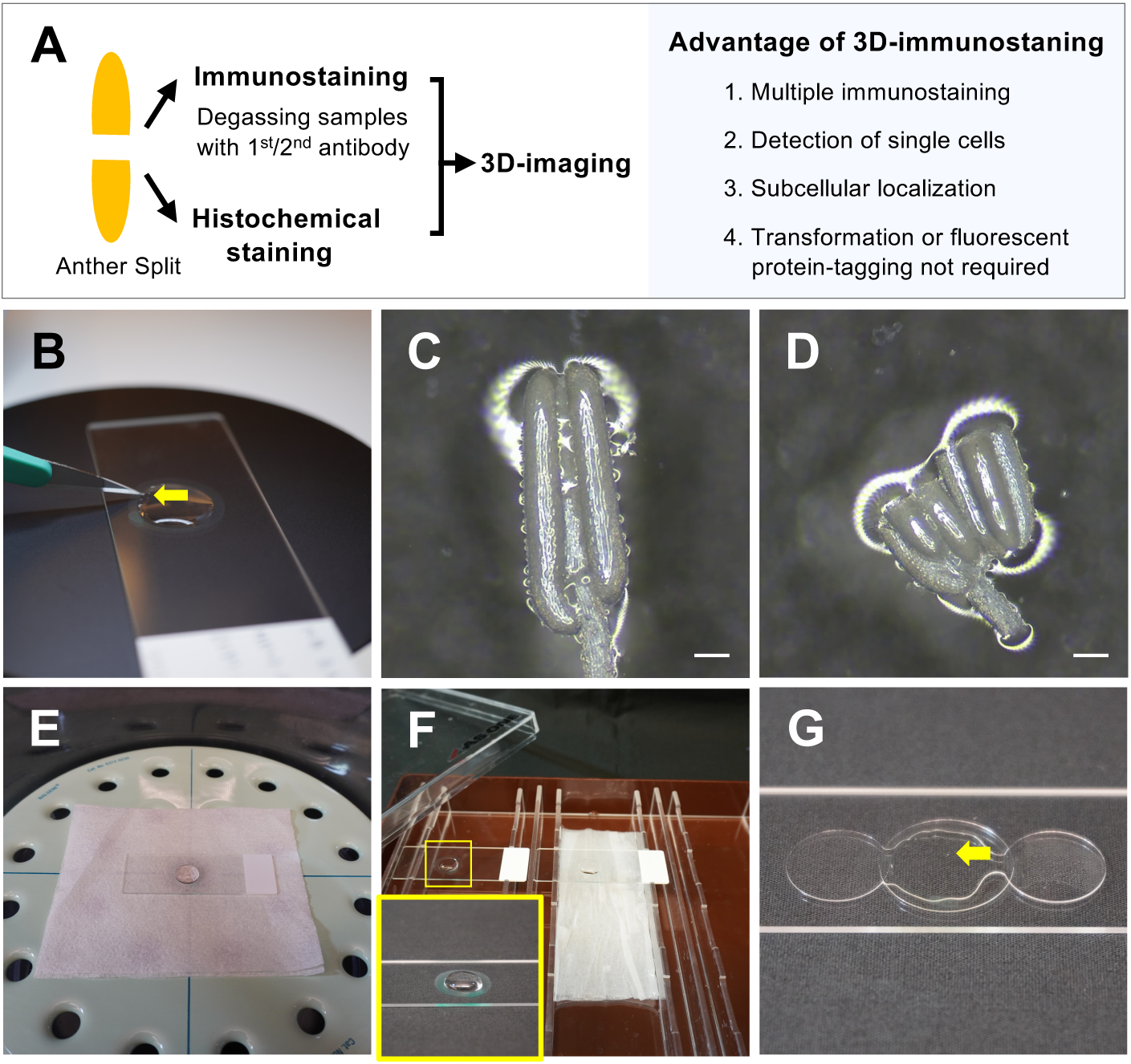
Equipment used for 3D-multiple immunostaining and split anthers. (**A**) Flow chart and advantages of the 3D-multiple immunostaining method. (**B**) Anthers split using a scalpel on the MAS-coated slide under the stereomicroscope (Nikon). (**C**) A 4% PFA-fixed anther in distilled water before the split. (**D**) The two parts of an anther after the split. Scale bars, 100 μm. (**E**) Samples with primary/secondary antibody solution in the vacuum desiccator. (**F**) Samples without coverslip placed in a humidified box for primary/secondary immunization. Yellow-lined square shows an enlarged image of the immunization on the slide. (**G**) Mounting of samples. Two coverslips, which are 0.13– 0.17 mm in thickness (MATSUNAMI Micro Cover Glass 13 mm No. 1), were placed on either side to maintain the structure of anthers without crushing. Yellow arrow indicates the split anthers.

After the split and blocking steps, the anthers were placed in primary MEL1 and ZEP1 antibody solutions, 1/500 with 3% bovine serum albumin (BSA) in PME buffer (as mentioned in more detail in methods). Next, degassing for the primary and secondary immunostaining steps is required for antibody penetration into the whole anthers. In this step, the slides with antibodies solution were placed in the vacuum desiccator (NALGENE), degassed at 0.05 MPa for 2 min, and this step was repeated four more times (Fig. 2E). Then, the slides without the coverslip on the samples were transferred to a humidified box and incubated overnight at 4 °C (Fig. 2F). After washing, samples were stained with 4’,6-diamidino-2-phenylindole (DAPI) and mounted in ProLong Gold; the two coverslips (Matsunami Micro Cover Glass No. 1 0.13–0.17 mm) were placed on either side of the samples to avoid crushing the anther locules (Fig. 2G).

### Visualization of the 3D-anther immunostaining

We captured the images using an LSM 880 microscope (Carl Zeiss) and created images using ZEN or Imaris 9 (Movie 1; Fig. 3). It is possible to distinguish the four somatic cell layers in both “z” section, which refers to the longitudinal sections of anthers (Fig. 3B) and the “x” section, which indicates the cross-sections (Fig. 3A). DAPI signals were also detected in all somatic anther walls, the epidermis (Ep), endothecium (En), middle layer (Ml), and tapetum (Ta). Moreover, the localization of MEL1 and ZEP1 was restricted to the meiocytes. In particular, MEL1 was detected in the cytoplasm, while ZEP1 was identified on the meiotic chromosomes, indicating that the 3D immunostaining is a multiplex imaging method that allows for differentiating between the four somatic cell layers and the germ cell, and the subcellular localization of multiple proteins.

**Figure 3.**
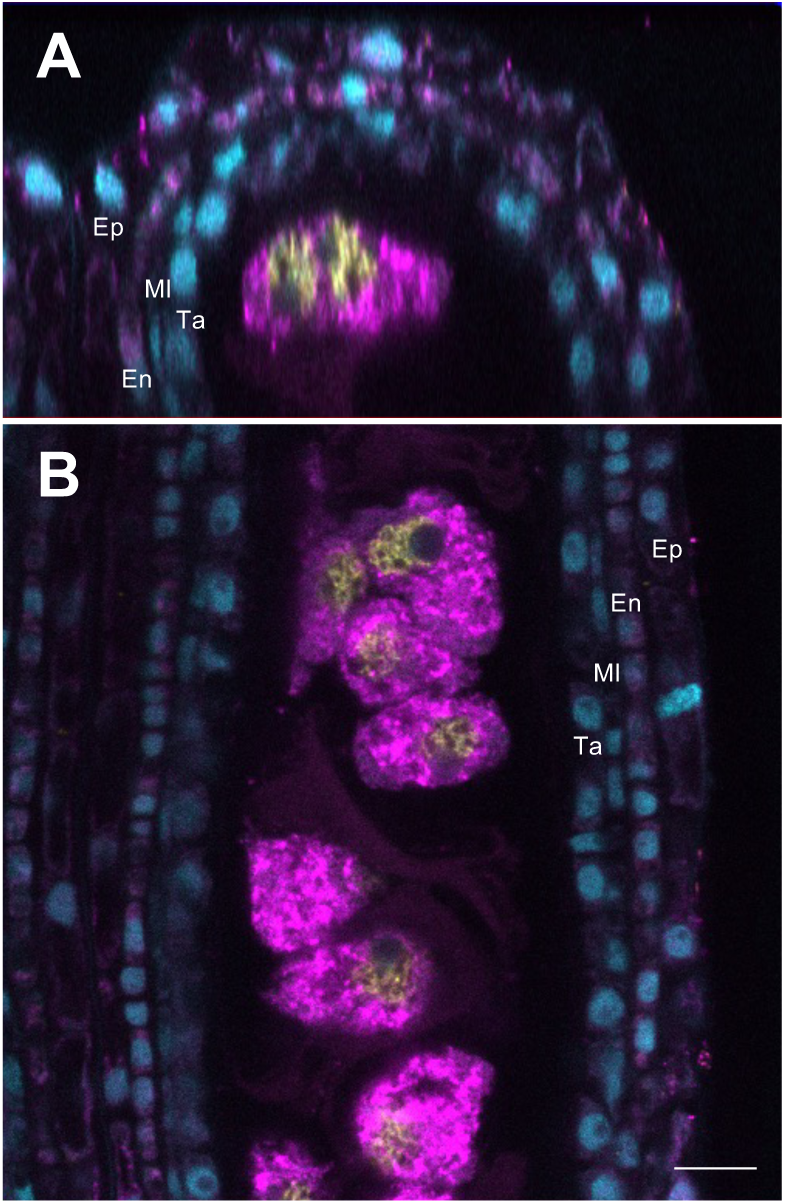
Cross and longitudinal sections of the 3D-anther immunoimaging using a confocal microscope. (**A**) The cross-section (X section) of the 3D-anther immunoimaging, in which 0.65 mm anthers from the early meiosis stage were used. (**B**) The longitudinal section (Z section) of the 3D-anther immunoimaging. The four somatic layers, epidermis (Ep), endothecium (En), middle layer (Ml), and tapetum (Ta), were observed. The cytoplasmic localization of MEL1 (magenta) and ZEP1 fluorescence signals (yellow) were observed in meiocytes surrounded by somatic anther walls. Cyan signals indicate DAPI staining. Scale bar, 10 μm. Laser excitation/emission are 405 nm/410–483 nm for DAPI, 561 nm/571–633 nm for MEL1, and 633 nm/638–755 nm for ZEP1.

Next, we investigated the 3D-histochemical staining using whole anthers. We also split the 4% fixative anthers, which are 0.5 mm long in the early meiosis stage, into two parts, and stained them with propidium iodide (PI) to stain DNA/RNA. We detected fluorescence signals in four anther walls and pollen mother cells (Supplementary Fig. S2). Consequently, these methods using the split anthers are effective for 3D-immunostaining and 3D-histochemical staining (Fig. 2A).

### Single-cell structures and MEL1/ZEP1 subcellular localizations using the whole-anther immunostaining method

Next, we explored the structure of single cells in each cell layer and in meiocytes of rice anthers during early meiosis using the 3D-multiple immunoimaging method. The epidermis cells were the largest in the anther wall layers (Fig. 4 A-D), and the endothecium cells exhibited elongated-transverse cell shapes (Fig. 4 E-H). The nucleus of the middle layer showed a characteristic structure (not round), whereas the nucleus in the tapetum layers was round (Fig. 4 I-P). MEL1 was primarily present in the cytoplasm in meiocytes (Fig. 4 S, T). In contrast to the MEL1-cytoplasmic localization, ZEP1 was detected in the nucleus region (Fig. 4 R, T). Notably, the filamentous signals of ZEP1 were detected on the meiotic chromosomes, which coincide with the DAPI-stained meiotic chromosomes (Fig. 4 Q, R, T). The different intercellular localization of the cytoplasmic MEL1 and nuclear-filamentous ZEP1 demonstrates that the simple 3D-multiple immunoimaging distinguishes the subcellular localization in each single cell of anthers.

**Figure 4.**
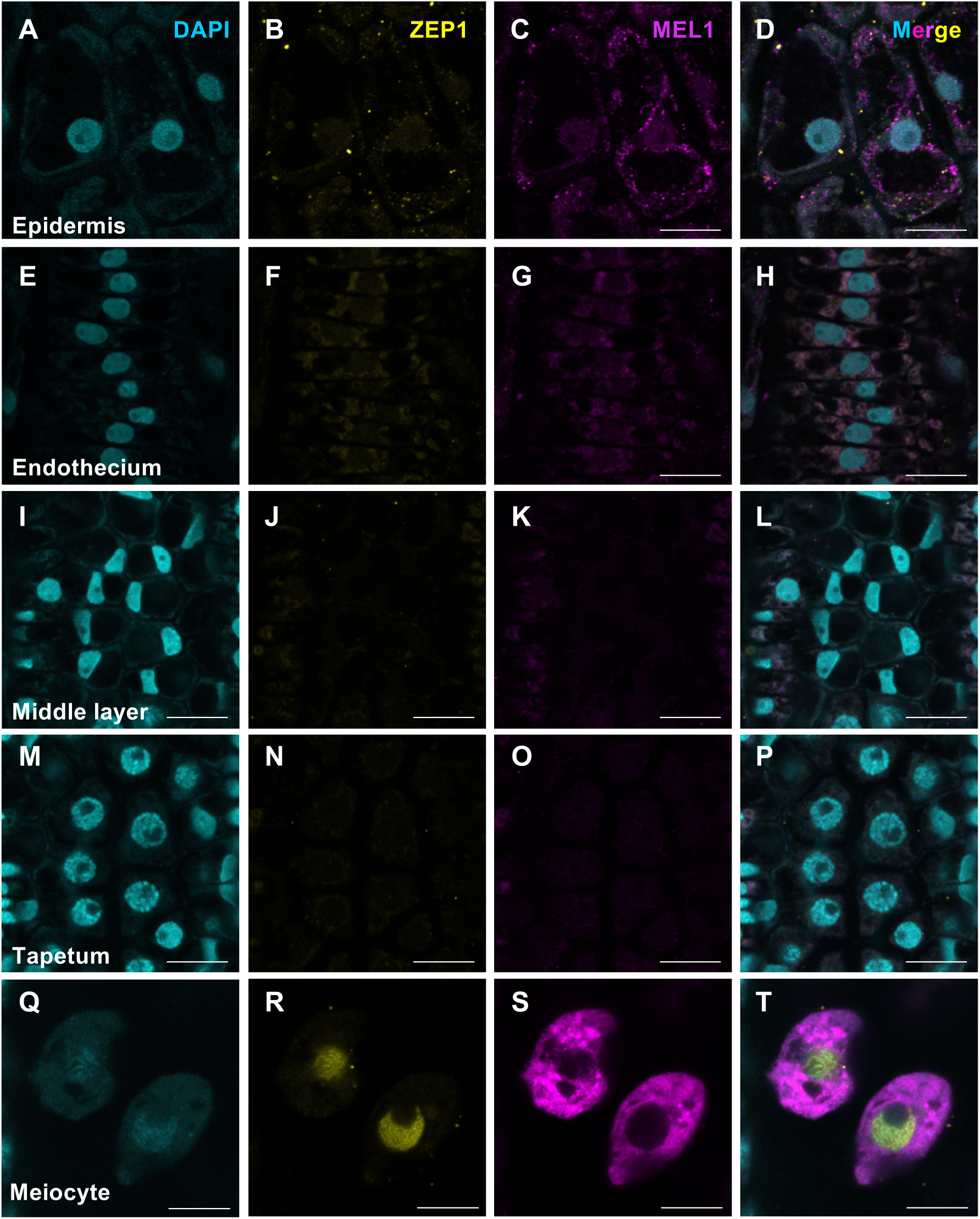
Single-cell structures and MEL1/ZEP1 subcellular localizations using the whole-anther immunostaining method. (**AEIMQ**) DAPI signals detected (cyan) in 3D-multiple immunoimaging using 0.65 mm-long whole anthers at early meiosis. (**B, F, J, N, and R**) 3D-multiple immunoimaging using against ZEP1(Yellow). (**C, G, K, O, and S)** 3D-multiple immunoimaging using against MEL1 (Magenta). (**D, H, L, P, and T**) Merged images of A-C, E-G, I-K, M-O, and Q-S, respectively. This 3D-immunostaining method enables in distinguishing cell types, the epidermis (Ep, **A-D**), endothecium (En, **E-H**), middle layer (Ml, **I-L**), tapetum layer (Ta, **M-P**), and meiocytes (**Q-T**). Scale bars, 10 μm. Laser excitation/emission are 405 nm/410–483 nm for DAPI, 561 nm/571–633 nm for MEL1, and 633 nm/638–755 nm for ZEP1.

## Discussion

In this study, we developed a 3D-imaging method that included histochemical staining and multiple immunostaining, for the study of whole anthers, which enabled us to distinguish between single cells out of the four somatic cell types in the anther without sectioning the anthers. Furthermore, it was possible to clearly detect the filamentous signals of ZEP1 on the meiotic chromosomes in addition to the specific subcellular localization of MEL1/ZEP1, indicating that 3D-immunoimaging methods could clarify the organelle localization in each single cell.

iTOMEI clearing is a powerful method to reduce the autofluorescence of plant organs ^13,18^. It was also applicable for the 3D-histochemical staining with PI in this study (Supplementary Fig. S2). However, we performed the iTOMEI clearing after 3D-immunoimaging of anthers, and it was difficult to detect the fluorescence signals, suggesting that the clearing method is not suitable for immunostaining using Alexa Fluor 567 and Alexa Fluor 678. Because the autofluorescence of anthers is considerably weaker than that of leaves, the clearing step is not required for 3D-anther immunoimaging. Using this simple-immunostaining method, it was possible to observe the fluorescence signals of the middle of an anther locule, wherein the depth is 50 μm, highlighting the detection of all types of somatic anther walls and germ in anthers without clearing.

One of the advantages of this immunoimaging method is that it does not require the transformation step to generate plants with fluorescence proteins. There is also no need for resin embedding, sectioning of samples or clearing during 3D-multiple immunoimaging (Fig. 2A). The steps of anther splitting and degassing prior to immunization are important for successful 3D-multiple immunostaining of anthers. The all procedures are performed on slide, not using the slide lack and staining jar for treatment and washing. Moreover, multiple immunoimaging could be performed using several antibodies to elucidate the localization of more than three proteins simultaneously and can be used at high resolutions to investigate subcellular localization in single cells. Improving this 3D-immunoimaging system would lead to the development of 3D-fluorescence *in situ* hybridization (3D-FISH) using the whole anther to detect the subcellular localization of mRNAs, long non-coding RNAs, and small RNAs in the near future. A large number of 21- and 24-nucleotide small RNAs are produced specifically in the anther and are loaded into the reproductive AGOs ^19,20^. It has also been reported that 24-nt small RNAs of the anther move from the tapetum to the meiocytes ^21^, indicating the possibility of non-cell-autonomous control via small RNAs as mobile signals in anthers. Hence, the method will also have potential applicability in observing the multiple localization of proteins and RNAs. The developed 3D-immunostaining method is a significant contribution to the literature because organ-level elucidation of the comprehensive molecular mechanisms underlying cell-to-cell interactions can be effectively applied to other living systems.

## Methods

### Plant materials and growth conditions

The *Japonica* rice ‘Nipponbare’ was used in this study. Plants were grown in growth chambers at 70% humidity with daily cycles of 14 h of light at 29.5–30 °C and 10 h of darkness at 25 °C for 40 days and then transferred to the short-day conditions with 10 h of light and 14 h of darkness, to promote the induction of the reproductive stage and to adjust the sampling stages.

### MEL1 and ZEP1 antibody generation

Two oligopeptides, MEL1-1 (CVYGAPMPAAHHQGAYQ) and MEL1-2 (GQAVAREGPVEVRQLPKC), were used to raise antibodies in rabbits (Cosmo Bio Co., LTD). The rabbit antiserum was purified using affinity chromatography (Cosmo Bio Co., LTD).

An N-terminal region of ZEP1 cDNA encoding was amplified using PCR using the following primer pair: 5’-gacaagcttgcggccATGCAGAAGCTGGGTTTATC-3’ and 5’-tgctcgagtgcggcctgTTCAGCAGATCTAGAATCCTCC-3’. The lowercase bases indicate the sequences of the pET24(+) vector for infusion. The ZEP1 amplicon was cloned into pET24(+) (Novagen, 69772) using an infusion system. The transformation to BL21 (DE3) pLys, purification, and immunization of rat were performed (SCRUM Inc.).

### Wes

Total proteins were extracted from 0.6–0.7-mm long anthers. The anthers were ground and mixed with an extraction buffer (150 mM NaCl, 50 mM Tris–HCl (pH 7.5), 0.1 % Tween 20, 10 % glycerol, 5 mM DTT, 1 mM Pefabloc SC (Roche), 1X Complete Protease Inhibitor Cocktail). After two rounds of centrifugation (5,800*g* for 10 min at 4 °C and 20,400*g* for 10 min at 4 °C) and removal of the debris, total proteins were extracted. MEL1 and ZEP1 antibodies (1/20 dilution) were used for the initial immune reactions.

### 4% paraformaldehyde fixation

Fixation of anthers was performed as previously described ^11^. The PFA fixative was prepared fresh immediately before use, and 4% PFA (Alfa Aesar) was added in 1X PMEG buffer (50 mM PIPES (Dojindo Molecular Technologies), 10 mM EGTA, 5 mM MgSO^4^·7H^2^O, 4% glycerol, 0.2% DMSO; pH 6.8). The inflorescences in the PFA fixative were degassed at 0.09–0.1 MPa for 20 min on ice (Supplementary Fig. S3), and this step was repeated three more times. The samples were incubated for 100 min at 25 °C and washed in 1X PMEG buffer for 20 min at 25 °C; this wash step was repeated five more times. The fixed inflorescences can be stored at 4 °C for six months.

### Immunostaining using pollen mother cells for antibody estimation

To release the meiocytes from anthers, the anthers from fixed inflorescence were incubated in an enzyme cocktail which contained 2% cellulase Onozuka-RS (Yakult Honsha, Japan), 0.3% pectolyase Y-23 (Kikkoman), and 0.5% Macerozyme-R10 (Yakult Honsha) in PME buffer (50 mM PIPES, 5 mM EGTA, and 5 mM MgSO^4^, pH 6.9) on a MAS-coated microscope slide, for 1 minute at 25°C. The anthers were washed with PME, squashed in distilled water using a needle to release meiocytes, and incubated at 25 °C for 30 minutes. The meiocytes were then blocked with 3% BSA in PME for 60 minutes. The meiocytes were incubated at 4 °C overnight with rabbit anti-MEL1 antibody or rat anti-ZEP1, diluted 1/500 with 3% BSA in PME. After washing three times with PME for 5 minutes, the slide was incubated in a dark chamber for 3 h at 25 °C with Goat anti-Rabbit IgG (H+L) Highly Cross-Adsorbed Secondary Antibody, Alexa Fluor 488 (Invitrogen, A11034) and Goat anti-Rat IgG (H+L) Cross-Adsorbed Secondary Antibody, Alexa Fluor 647 (Invitrogen, A21247), diluted 1/200 with 3% BSA/PME, followed by three washes with PME for 5 minutes each. Then, the MEL1 sample was mounted in the Vectashield mounting medium with DAPI (Vector Laboratories, H-1200), and the image was captured using an LSM780 microscope (Carl Zeiss). ZEP1 samples were incubated for 15 min at 25 °C in DAPI (SIGMA, MBD0015), followed by washing three times with PME buffer. Samples were mounted in the ProLong Gold antifade reagent (Invitrogen, P10144) and the image was captured using an LSM880 microscope (Carl Zeiss).

### Immunostaining using a whole mount of anthers

The anthers were retrieved from 4% paraformaldehyde-fixed inflorescences in PME buffer under a stereomicroscope. Next, the anthers were transferred to distilled water in a circle drawn with a PAP pen (Daido Sangyo) on a MAS-coated microscope slide (Matsunami Glass) and split into two parts using a scalpel; the samples were incubated for 30 minutes at 25 °C. Then, the cut anthers were blocked with 3% BSA in PME for 60 minutes at 25 °C. The samples were placed in primary antibody solutions (Rabbit anti-MEL1, rat anti-ZEP1 diluted 1/500 with 3% BSA in PME), degassed at 0.05 MPa for 2 minutes at 25 °C, which was repeated four more times, and incubated overnight at 4 °C. After washing three times with PME for 5 minutes, the slide was placed in secondary antibody solutions, anti-Rabbit IgG, Alexa Fluor 568 (Invitrogen, A11036) and anti-Rat IgG, Alexa Fluor 647 (Invitrogen, A21247) diluted 1/200 with 3% BSA in PME, degassed at 0.05 MPa for 2 minutes at 25 °C, which was repeated four more times. The slide was incubated in a dark humid chamber for 2 h at 25 °C and then incubated overnight at 4 °C. The samples were washed three times with PME buffer for 5 minutes. The samples were then incubated for 15 minutes at 25 °C in DAPI (SIGMA, MBD0015), followed by three washes with PME buffer. Samples were mounted in the ProLong Gold antifade reagent (Invitrogen, P10144) with a coverslip (Matsunami Glass).

### Visualization of the 3D immunostaining of the entire anthers

The images was captured using an LSM880 microscope (Carl Zeiss). Conditions: 40x (1.3 oil) Plan Apochromat lens (Fig. 3 and Movie1) and 63x (1.4 oil) Plan Apochromat lens (Fig. 4) for detection, 405, 561, and 633 nm laser lines for DAPI, Alexa Fluor 568, and Alexa Fluor 647 excitation, 410–483 nm (DAPI), 571–633 nm (Alexa Fluor 568) and 638–755 nm (Alexa Fluor 647) filter emission. The images and animation were created using the ZEN (Carl Zeiss) or Imaris 9 (Bitplane AG) software (Figs. 3, 4; Movie 1).

### 3D-histochemical staining of whole anthers and imaging

Anthers fixed in 4% paraformaldehyde were split into two parts using a scalpel. The anthers were stained with PI for 15 min at 25 °C. After PI staining, the samples were transferred to 20% iTOMEI solution (20% caprylyl sulfobetaine in 100 mM sodium phosphate buffer) and incubated for 10 minutes at 25 °C. The samples were then transferred and incubated in 50% iTOMEI solution (50% caprylyl sulfobetaine in 100 mM sodium phosphate buffer) for 10 minutes at 25 °C. Finally, the samples were transferred and incubated in 70.4% iTOMEI (70.4% iohexol in PBS) for 1 h at 25 °C ^13^. The samples were mounted with 70.4% iTOMEI. Images were captured with a confocal microscope (LSM880; Carl Zeiss). Conditions: 40x (1.3 oil) Plan Apochromat lens for detection, 561 nm laser lines for PI excitation, 571–633 nm filter emission. The images were created using ZEN (Carl Zeiss) (Supplementary Fig. S1).

## Acknowledgements

This work was supported by the JST FORESTO Program (Grant Number JPMJFR204U, Japan), JST PRESTO Program (Grant Number JPMJPR17Q3, Japan), the Naito Foundation, and the Okinawa Institute of Science and Technology Graduate University, Japan. We thank Science and Technology Group members and FORESTO members (Siomi panel) for their helpful discussions.

## Author contributions

R.K. conceived the study, conducted most of the data analysis, and drafted the manuscript.

S.A. and R.K. performed 3D immunostaining, and H.T. performed protein experiments.

S.A. assisted in creating imaging figures.

## Movie and figure legends

**Movie 1.**
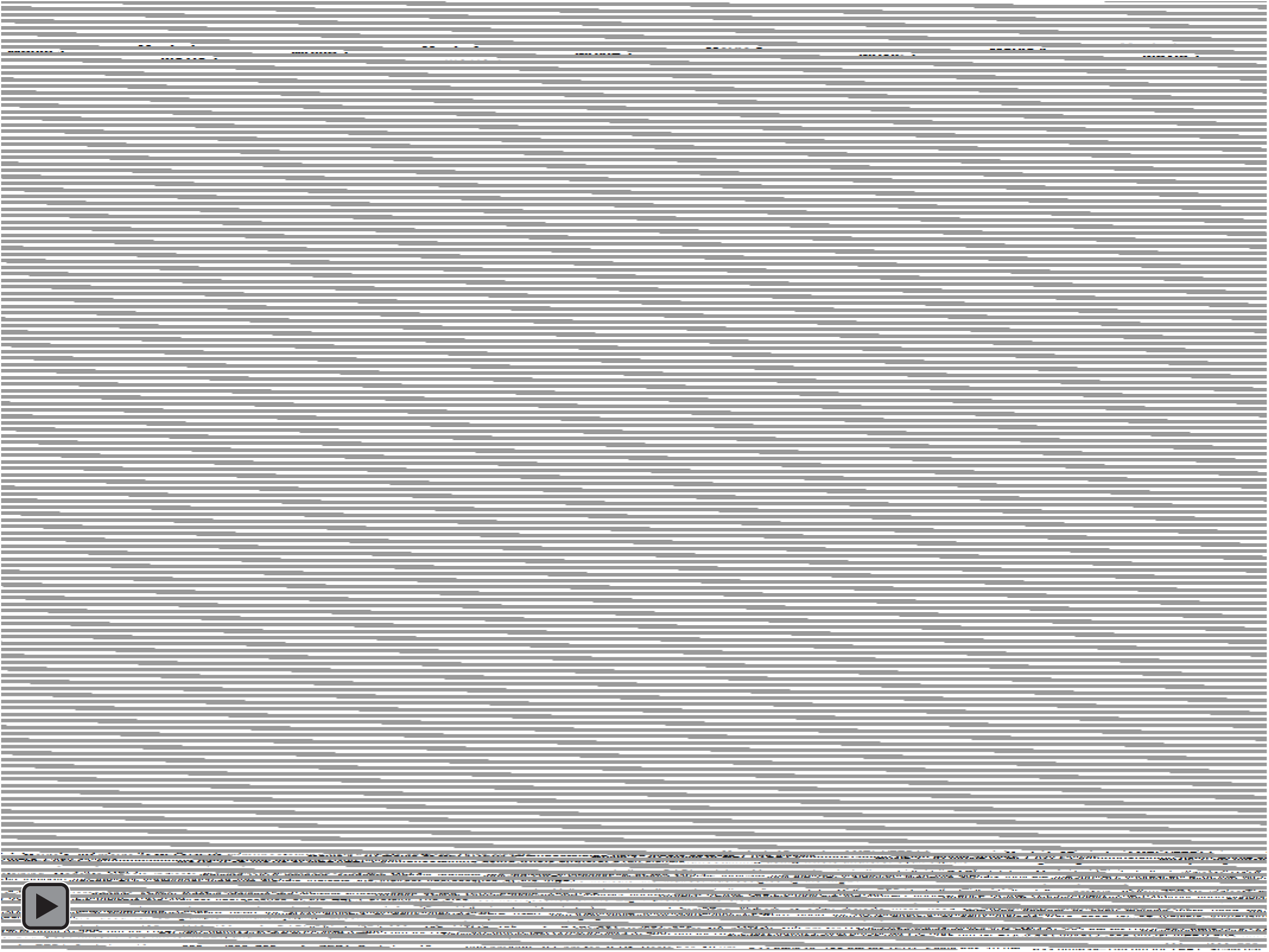
3D movie of MEL1/ZEP1 immunostaining using whole anthers. Cyan signals indicate DAPI staining. Magenta signals indicate the indirect fluorescence of the MEL1 protein. Yellow signals indicate the indirect fluorescence of the ZEP1 protein. The 0.65 mm-long anthers at early meiosis were used for 3D-multiple immunoimaging. Laser excitation/emission are 405 nm/410–483 nm for DAPI, 561 nm/571–633 nm for MEL1, and 633 nm/638–755 nm for ZEP1. Scale bar, 10 μm.

**Figure S1.**
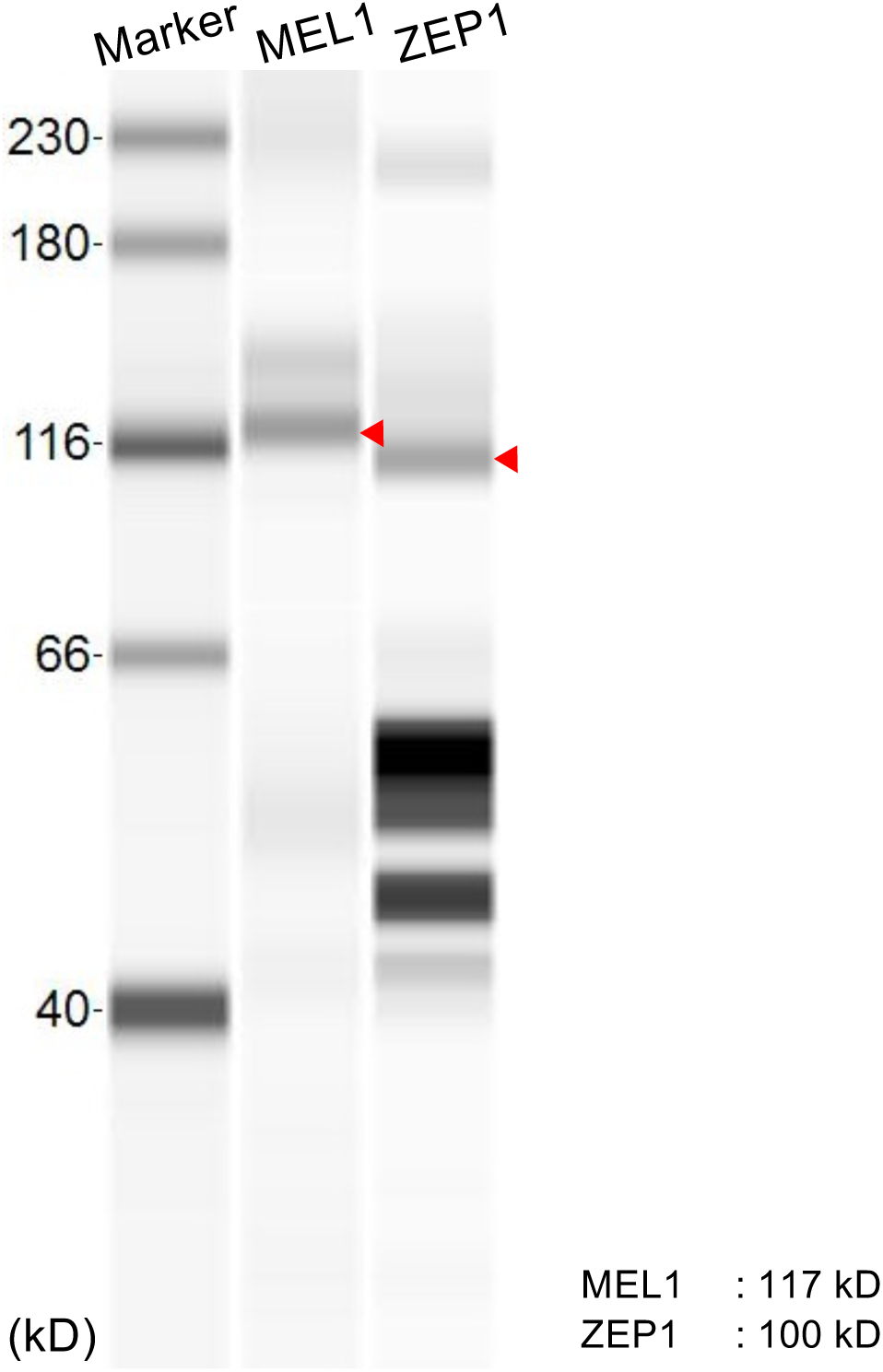
Wes analysis of MEL1 and ZEP1 proteins in anthers during early meiosis. Total proteins were extracted from 0.6–0.7 mm of anthers. MEL1 and ZEP1 proteins were enriched during early meiosis. Wes signals for each MEL1/ZEP1 proteins coincide with the predicted molecular weights, demonstrating that the anti-MEL1 or anti-ZEP1 antibodies detect MEL1/ZEP1 proteins. Red arrows indicate the molecular weight of the MEL1 and ZEP1 proteins.

**Figure S2.**
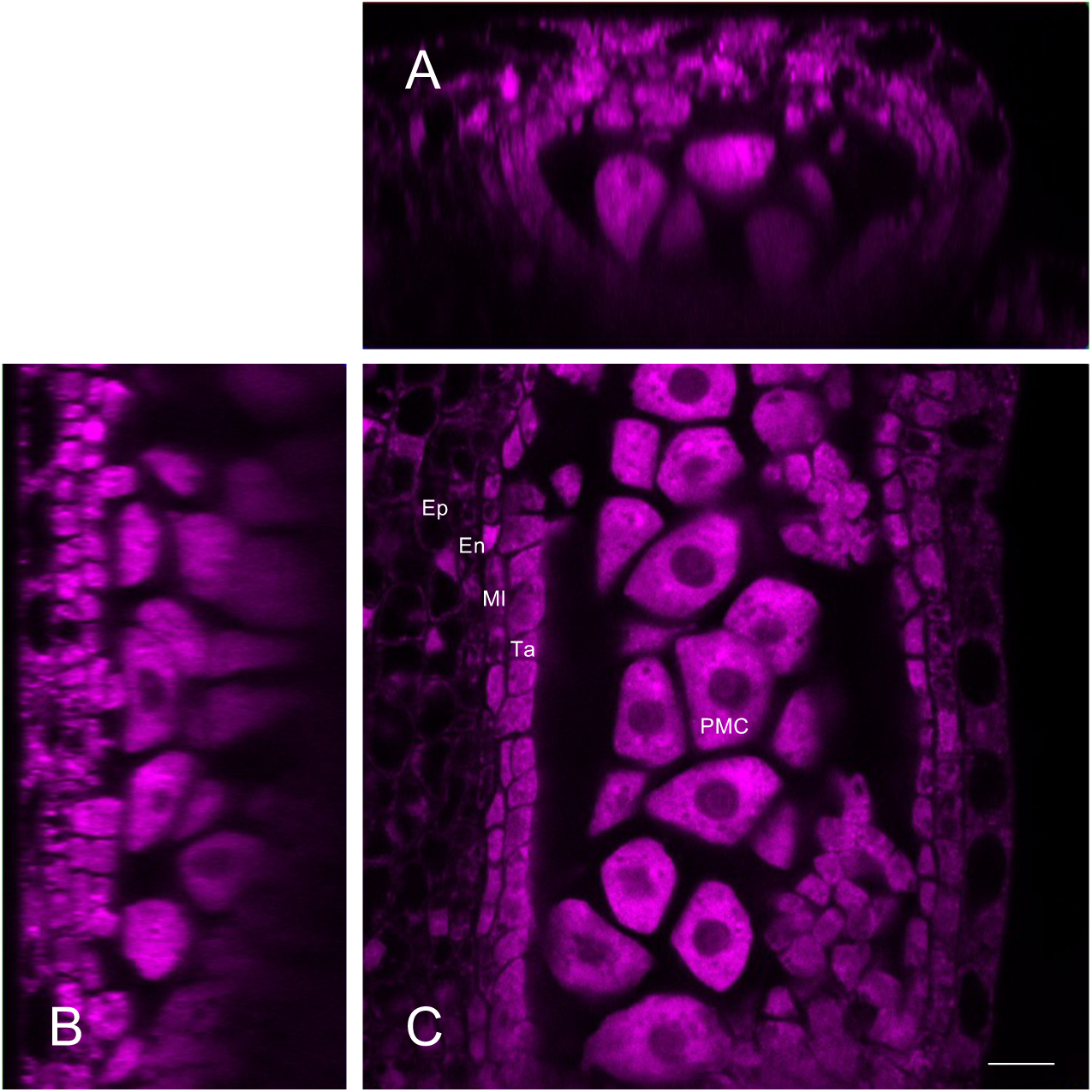
3D-histochemical imaging using whole anthers. (**A**) The cross-section (x section) of the PI-staining imaging of the 0.5 mm-long anthers. (**B**) The y section of the PI-staining imaging of the 0.5 mm-long anthers. (**C**) The longitudinal section (z section) of the PI-staining imaging of the 0.5 mm-long anthers. The four somatic layers, the epidermis (Ep), endothecium (En), middle layer (Ml), and tapetum (Ta), and pollen mother cells (PMC) were observed. Scale bar, 10 μm.

**Figure S3.**
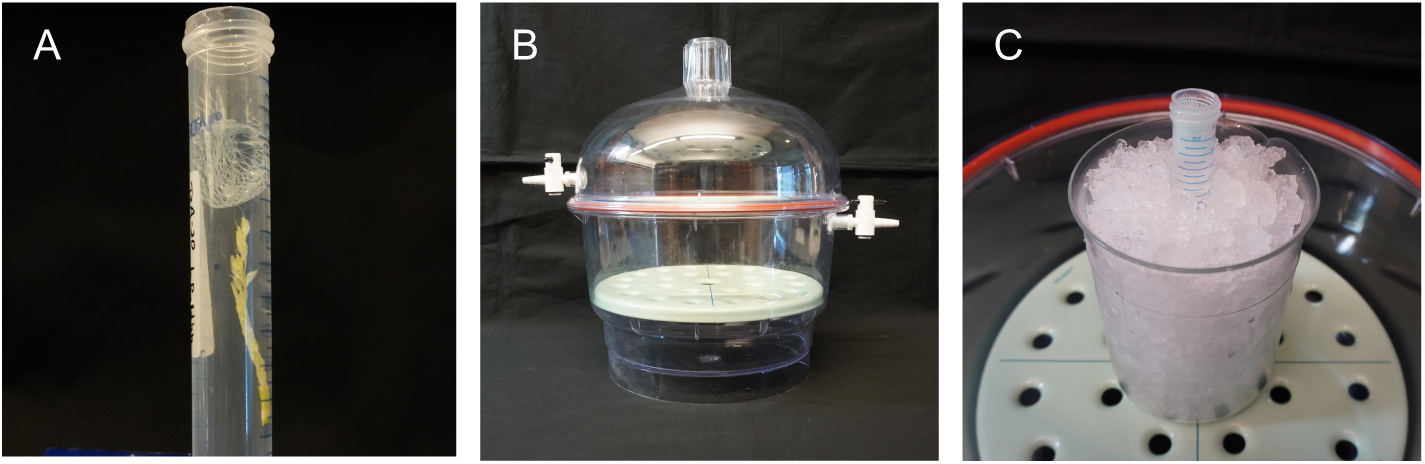
Equipment used for anther fixation. (**A**) Inflorescences in 4% PFA fixative using a 15 mL plastic tube. (**B**) The vacuum desiccator for the fixation. (**C**) the 15 mL plastic tube with samples on ice during the fixation.

**Supplementary Table S1.** Developmental stages of anthers.

| Stages | Anther (mm) | Inflorescence (mm) | Germ cell development | Anther wall development | MEL1 localization | ZEP1 localization |
| --- | --- | --- | --- | --- | --- | --- |
| stage 1 | 0.3-0.4 | 2.0 | Premeiosis<br>(Primordial germ cell initiation) | 4 layers<br>(Ep, En, MI, Ta) | Cytoplasm | - |
| stage 2 | 0.5-0.7 | 3.0-4.0 | Leptotene--Diplotene | Ta differentiation | Cytoplasm | filamentous signals<br>on meiotic<br>chromosomes |
| stage 3 | 0.6-0.7 | 3.0-5.0 | Meiotic Division | Ta differentiation | - | - |
| stage 4 | 0.8-0.9 | 5.0-6.0 | Microspore | Ta programmed cell death<br>MI disappearance | N.D. | N.D. |

Ep:Epidermis, En:Endothecium, MI:Middle layer, Ta:Tapetum, "-": no observation, N.D.: no data

## References

1 Malone, C. D. et al. Specialized piRNA pathways act in germline and somatic tissues of the Drosophila ovary. Cell 137, 522–535, doi:10.1016/j.cell.2009.03.040 (2009).

2 Zhai, J. et al. Spatiotemporally dynamic, cell-type-dependent premeiotic and meiotic phasiRNAs in maize anthers. Proc Natl Acad Sci U S A 112, 3146–3151, doi:10.1073/pnas.1418918112 (2015).

3 Komiya, R. Biogenesis of diverse plant phasiRNAs involves an miRNA-trigger and Dicer-processing. J Plant Res 130, 17–23, doi:10.1007/s10265-016-0878-0 (2017).

4 Iwasaki, Y. W., Siomi, M. C. & Siomi, H. PIWI-Interacting RNA: Its Biogenesis and Functions. Annu Rev Biochem 84, 405–433, doi:10.1146/annurev-biochem-060614-034258 (2015).

5 Komiya, R. Spatiotemporal regulation and roles of reproductive phasiRNAs in plants. Genes Genet Syst 96, 209–215, doi:10.1266/ggs.21-00042 (2022).

6 Nonomura, K. et al. The MSP1 gene is necessary to restrict the number of cells entering into male and female sporogenesis and to initiate anther wall formation in rice. Plant Cell 15, 1728–1739, doi:10.1105/tpc.012401 (2003).

7 Vernoud, V. et al. The HD-ZIP IV transcription factor OCL4 is necessary for trichome patterning and anther development in maize. Plant J 59, 883–894, doi:10.1111/j.1365-313X.2009.03916.x (2009).

8 Olmedo-Monfil, V. et al. Control of female gamete formation by a small RNA pathway in Arabidopsis. Nature 464, 628–632, doi:10.1038/nature08828 (2010).

9 Matsuo, Y., Arimura, S. & Tsutsumi, N. Distribution of cellulosic wall in the anthers of Arabidopsis during microsporogenesis. Plant Cell Rep 32, 1743–1750, doi:10.1007/s00299-013-1487-1 (2013).

10 Araki, S. et al. miR2118-dependent U-rich phasiRNA production in rice anther wall development. Nat Commun 11, 3115, doi:10.1038/s41467-020-16637-3 (2020).

11 Koizumi, K. & Komiya, R. 3D imaging and in situ hybridization for uncovering the functions of microRNA in rice anther. Methods Mol Biol, in press.

12 Palmer, W. M. et al. PEA-CLARITY: 3D molecular imaging of whole plant organs. Scientific Reports 5, 13492, doi:10.1038/srep13492 (2015).

13 Sakamoto, Y. et al. Improved clearing method contributes to deep imaging of plant organs. Commun Biol 5, 12, doi:10.1038/s42003-021-02955-9 (2022).

14 Nagaki, K., Yamaji, N. & Murata, M. ePro-ClearSee: a simple immunohistochemical method that does not require sectioning of plant samples. Sci Rep 7, 42203, doi:10.1038/srep42203 (2017).

15 Komiya, R. et al. Rice germline-specific Argonaute MEL1 protein binds to phasiRNAs generated from more than 700 lincRNAs. Plant J 78, 385–397, doi:10.1111/tpj.12483 (2014).

16 Wang, M. et al. The central element protein ZEP1 of the synaptonemal complex regulates the number of crossovers during meiosis in rice. Plant Cell 22, 417–430, doi:10.1105/tpc.109.070789 (2010).

17 Nonomura, K. et al. A germ cell specific gene of the ARGONAUTE family is essential for the progression of premeiotic mitosis and meiosis during sporogenesis in rice. Plant Cell 19, 2583–2594, doi:10.1105/tpc.107.053199 (2007).

18 Sato, M., Akashi, H., Sakamoto, Y., Matsunaga, S. & Tsuji, H. Whole-Tissue Three-Dimensional Imaging of Rice at Single-Cell Resolution. Int J Mol Sci 23, doi:10.3390/ijms23010040 (2021).

19 Johnson, C. et al. Clusters and superclusters of phased small RNAs in the developing inflorescence of rice. Genome Res 19, 1429–1440, doi:10.1101/gr.089854.108 (2009).

20 Liu, Y., Teng, C., Xia, R. & Meyers, B. C. PhasiRNAs in Plants: Their Biogenesis, Genic Sources, and Roles in Stress Responses, Development, and Reproduction. Plant Cell 32, 3059–3080, doi:10.1105/tpc.20.00335 (2020).

21 Zhou, X. et al. 24-nt phasiRNAs move from tapetal to meiotic cells in maize anthers. bioRxiv, 2021.2008.2024.457464, doi:10.1101/2021.08.24.457464 (2022).

